# ROBOCOV: An affordable open-source robotic platform for SARS-CoV-2 testing by RT-qPCR

**DOI:** 10.1101/2020.06.11.140285

**Authors:** José Luis Villanueva-Cañas, Eva Gonzalez-Roca, Aitor Gastaminza Unanue, Esther Titos, Miguel Julián Martínez Yoldi, Andrea Vergara Gómez, Joan Antón Puig Butillé

## Abstract

Current global pandemic due to the SARS-CoV-2 has struggled and pushed the limits of global health systems. Supply chain disruptions and scarce availability of commercial laboratory reagents have motivated worldwide actors to search for alternative workflows to cope with the demand.

We have used the OT-2 open-source liquid-handling robots (Opentrons, NY), RNA extraction and RT-qPCR reagents to set-up a reproducible workflow for RT-qPCR SARS-CoV-2 testing. We developed a framework with a template and several functions and classes that allow the creation of customized RT-qPCR automated circuits. As a proof of concept, we provide data obtained by a fully-functional tested code using the MagMax™ Pathogen RNA/DNA kit and the MagMax™ Viral/Pathogen II kit (Thermo Fisher Scientific, MA) on the Kingfisher™ Flex instrument (Thermo Fisher Scientific, MA). With these codes available is easy to create new stations or circuits from scratch, adapt existing ones to changes in the experimental protocol, or perform fine adjustments to fulfil special needs.

The affordability of this platform makes it accessible for most laboratories and hospitals with a bioinformatician, democratising automated SARS-CoV-2 PCR testing and increasing, temporarily or not, the capacity of carrying out tests. It also confers flexibility, as this platform is not dependant on any particular commercial kit and can be quickly adapted to protocol changes or other special needs.

## INTRODUCTION

The rapid spread of the coronavirus disease 2019 (COVID-19), caused by SARS-CoV-2^1^, has had a huge health and economic impact worldwide. Since the identification of the virus, molecular tests for its rapid detection have become essential^2^. Early diagnosis has been crucial to limit the spread of the virus and to manage public health resources. Currently, the Reverse Transcription quantitative Polymerase Chain Reaction (RT-qPCR) is the main molecular approach for COVID-19 testing ^3^.

Large-scale testing for COVID-19 is also crucial in the de-escalation of mitigation measures. In the most stricken countries, the shortage of laboratory supplies and limited testing capacity have restricted the use of RT-qPCR to few population groups such as patients with severe symptoms, high-risk patients or healthcare workers. Population screening is crucial in disease control measures such as contact tracing and isolation. This is particularly challenging considering that pre-symptomatic^4^ and asymptomatic transmission^5^ seems to play a key role in the infection dynamics. The use of SARS-CoV-2 RT-qPCR as a screening tool in high-risk settings, such as hospitals, long-term care facilities or nursing homes, is a valuable measure to promptly isolate cases.

There are several commercial platforms for automated high-throughput testing, such as the cobas 6800 from Roche^4^ or the Panther Fusion system from Hologic^7^. However, these solutions are expensive and not within reach for most laboratories. These platforms usually work with proprietary reagent cartridges specifically tailored to each platform and labware with few or none generic suppliers, putting a crimp in testing if supply chains break. Other liquid handling automated workstations from companies such as TECAN or Hamilton are high throughput platforms that can accommodate different protocols if they have the necessary modules. In response to COVID-19 they all have adapted extraction, qPCR, and ELISA protocols into their machines ^8,9^. These platforms are programmed through their own proprietary software and users are able to create their own scripts. Although they have certain flexibility, their high price is still a barrier to many laboratories.

The advent of the affordable OT-2 open-source liquid handling robots, developed by Opentrons, has opened the door to low-cost automated solutions^10^. The stations, initially conceived to perform simple pipetting tasks in research labs, can be concatenated to produce more complex workflows. The OT-2 stations have a limited number of actions compared to other commercial workflows. However, their affordability and being open-source at both hardware and software levels make them an attractive solution for molecular laboratories. Their openness confers them the ability to avoid limited supply chains for certain laboratory equipment, as it takes little time to define and test new laboratory equipment and include it in an already working protocol^11^. Open software is another advantage, as any development can be shared at once by other users. In addition, Python is the language code used to develop protocols which currently is the third most used language worldwide with an estimated community of developers of 8.5 Million^12^.

In this work, we provide a general template and base code that can be used to develop custom stations and complete circuits in OT-2 robots. By using this custom framework, we developed several functional stations to automate the SARS-CoV-2 RT-qPCR testing. We provide code for distinct circuits, useful to increase the testing capacity of established laboratories and provide freedom during acute shortages. These circuits might be an optimal solution for laboratories that currently do not have automated platforms. We are compelled to share our knowledge while we continue developing the platform so other organizations can benefit from it.

## METHODS AND FUNCTIONALITIES

### Nasopharyngeal swab lysates preparation

Nasopharyngeal swab samples were collected and immediately placed into a sterile tube, containing 2 mL of lysis buffer (2 M guanidinium thiocyanate, 2 mM dithiothreitol, 30 mM sodium citrate, and 1 % Triton X-100). Samples were vortexed and mixed 1:1 with cobas omni Lysis Reagent (Roche, Basel, Switzerland) for inactivation.

### Nucleic acid extraction and SARS-CoV-2 qPCR set up

Viral RNA was extracted from nasopharyngeal swab lysates using either the MagMax™ Pathogen RNA/DNA kit and the MagMax™ Viral/Pathogen II kit on the Kingfisher™ Flex Purification System (Thermo Fisher Scientific, Waltham, MA) according to manufacturer’s instructions with some modifications. Specifically, using the MagMax™ Pathogen RNA/DNA kit, no addition of MagMax™ Pathogen RNA/DNA Lysis Solution Master Mix was applied to the inactivated samples and 460 µl of lysates were directly transferred to the King Fisher™ deep-well sample plate. Subsequently, 260 µl of MagMax™ Pathogen RNA/DNA Bead Master Mix and 5 µl of the MS2 Phage Control (Thermo Fisher Scientific, Waltham, MA) were added to each well including a negative control sample containing 460 µl of PBS located at A1 position of the 96 deep-well plate. Sample plate was then loaded onto the KingFisher™ instrument and automated RNA extraction was performed using the BindIt™ protocol “MagMax_Pathogen_High_Volume”.

Viral RNA extraction using the MagMax™ Viral/Pathogen II kit was performed according to the protocol for “400 µl sample input with two wash steps” recently published by Thermo Fisher Scientific as a “Sample Prep Application Note” at its web site with some modifications. Briefly, no addition of proteinase K was applied to the inactivated samples and 400 µl of lysates were directly transferred to the King Fisher™ deep-well sample plate. Subsequently, 550 µl of Binding Bead Mix and 10 µl of the MS2 Phage Control (Thermo Fisher Scientific, Waltham, MA) were added to each well including a negative control sample containing 400 µl of PBS located at A1 position of the 96 deep-well plate. Sample plate was then loaded onto the KingFisher™ instrument and automated RNA extraction was performed using the BindIt™ protocol “MVP_2Wash_400_Flex” provided in Thermo Fisher’s web site with minor modifications. Specifically, the proteinase K digestion step was removed and plastic configuration was used as follows: three King Fisher™ deep-well plates (samples, wash1 and wash2), 2 King Fisher™ Standard plates (eluates and place for Tip Comb) and one Tip Comb. See Availability Section for more information.

### Code Features

The code developed so far provide features that improve the base capabilities of the OT-2 platform and can be used as a starting point to develop complete circuits.

#### General template

A code template eases the development of new protocols and the automation of some tasks. The template has been designed to work with a variable number of samples and include the features described below. It is structured in distinct steps, making it easier to understand the code and execute all the steps or only those that are necessary. This is useful for testing or resuming a run following a problem.

#### Reagent classes

The liquids used in a protocol might have distinct physical properties. We developed a system that defines the liquids as objects with properties at the beginning of a protocol. With this system, it is only necessary to specify which liquid is being used in a step to automatically apply all the adjustments for that liquid. For example, when dispensing alcohol-based substances, a coating of the liquid is left on the tip; in these scenarios the attribute “rinse” can be enabled. Pre-rinse is a simple technique to increase pipetting accuracy through the pre-conditioning of tips by aspirating a certain volume prior to using the tip. A more detailed description of this function is available on GitHub.

#### Height adjustments

The reagent classes store the total volumes, location, and remaining volumes of all liquids defined. By using this information combined with the geometrical measures of the containers, we developed a simple formula to calculate the optimal aspiration height when pipetting that liquid.

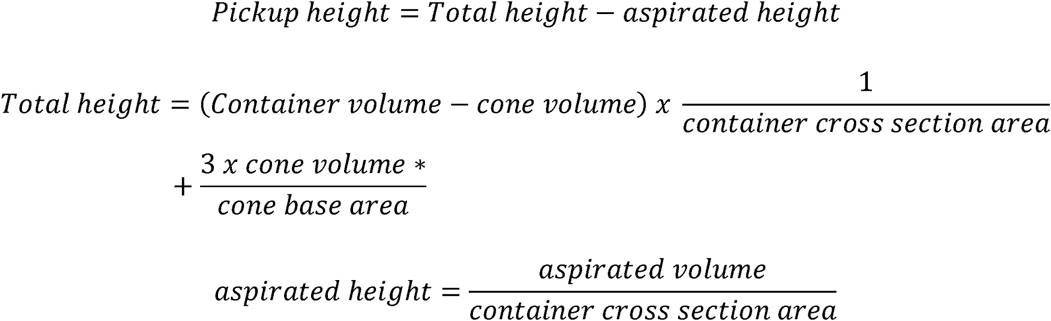

*Formula for a container with cone shaped bottom. The formula should be adjusted for a different shape. See GitHub for details.

#### Other enhanced functions

These include a highly customizable liquid transfer function, which makes use of the reagent classes to adjust different parameters, a multi-dispensing function for one to many transfers or a reagent mixing function, among others.

#### Log system

Currently, every run generates a time report, documenting how much time every step took as well as reporting the number of tips used in a run. These reports are .txt files that are stored in the robot and can be easily retrieved if necessary.

#### Automation

An important aspect of implementing a circuit in clinical diagnostics is the traceability of samples. This is difficult to generalize as each laboratory has its own protocols, LIMS, and databases. We provide several scripts general enough to be adapted to different working environments. For example, a template for sample barcode scanning that tracks sample placement in the different possible dispositions (i.e. 4 racks with 24 samples each, qPCR with 96 wells, etc.) or the automatic generation of templates with sample names and well positions for the ABI 7500 qPCR system (Thermo Fisher Scientific) or the LightCycler 480 II system (Roche life science).

#### Custom labware

Definitions of commonly used plasticware from different companies are provided. They are available as .json files with the measures and their description and can be imported into the OT-2 app.

### Interpretation of SARS-CoV-2 qPCR results

The results from the ABI 7500 qPCR system generated by the 7500 software v2.3 (Thermo Fisher Scientific) are exported as .csv files and later transformed into a user-friendly table using an Rmarkdown script, including an automatic interpretation of results. A sample is considered negative when the internal control is amplified but the viral targets are not, invalid when there is no amplification of the internal control and positive when two or more targets are amplified. Other cases are marked as “revise” and individually evaluated prior to transferring the results to the laboratory information system.

## RESULTS

### Calculation of aspiration height of the OT-2 pipettes

A common technique to manually pipette is to plunge the tip a few mm below the meniscus. We risk aspirating air if we plunge too little and too much immersion can cause liquid to cling to the outside of the tip. However, OT-2 stations do not have conductive tips to detect a first contact with liquid. To amend this problem, we developed a function that calculates the most suitable aspiration height for the pipette based on the remaining volume available and the shape of the container (see Methods). It also considers that a reagent can be in multiple sources and switches to another source if there is not enough volume in the current one (i.e. different screwcaps). This is useful to increase volume accuracy, to avoid overspills, and to efficiently use limiting reagents. In Figure 1 we show a simulation of a common scenario where we have two screwcaps with a limiting reagent. Initial volumes can be adjusted using simulations to minimize reagent leftover.

**Figure 1.**
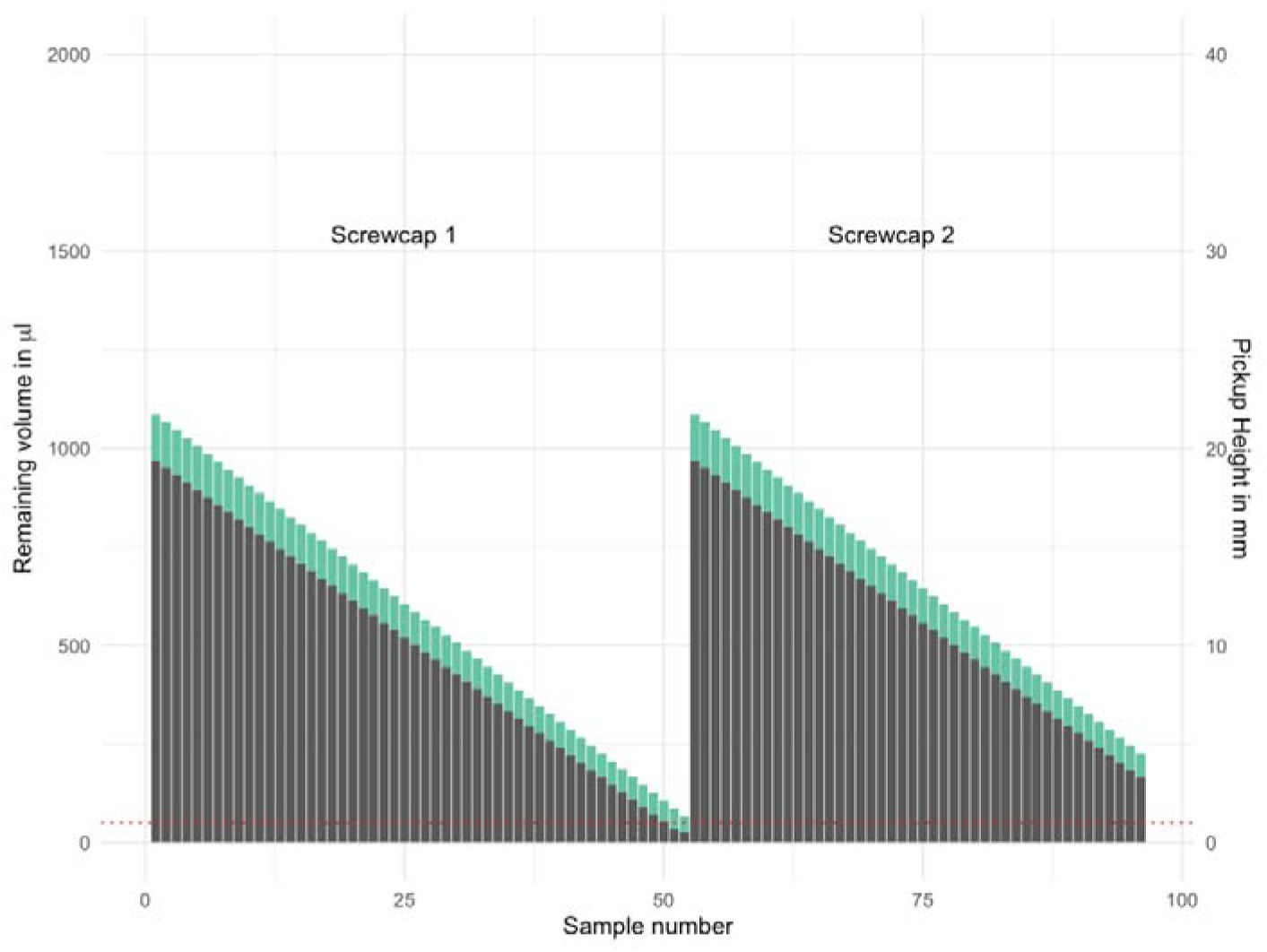
Simulations using the height calculation function. The plot represents two 2 ml screwcaps with a reagent that is used in 96 wells (one full plate), where 20 µl are dispensed in each one. The initial reagent is2112 µl and includes an initial 10% extra volume (20 µl * 96 samples * 1.1) to compensate for pipette inaccuracies. The left y axis (green) represents the remaining volume in each iteration after aspirating 20 µl. The right x axis (dark grey) indicates the height from where we are aspirating. The red dotted line depicts the volume that fits in the screwcap cone (50 µl). If there is less volume than the needed aspirated volume (20 µl) the system will switch to the next reagent location (screwcap 2).

### OT-2/Kingfisher circuits: OT-2-KF pathogen and OT-2 KF viral pathogen II

We have built and tested different functional circuits using our code framework in the OT-2 stations. As a proof of concept, we provide the code and results for two circuits that work with equipments and reagents from Thermo Fisher Scientific, which are currently used for SARS-CoV-2 testing in our laboratory. The circuits are named KF pathogen and KF viral pathogen II, after the extraction kits used.

A circuit is composed by six stations: four OT-2 stations, a Kingfisher™ Flex Purification System, and one ABI 7500 Fast Real-time PCR instrument. The difference between both circuits relies on the kit used for RNA extraction: MagMax™ Pathogen RNA/DNA kit or MagMax™ Viral Pathogen II RNA/DNA kit Both circuits use TaqPath™ COVID-19 CE-IVD RT-PCR reagents for the RT-qPCR reaction.

The circuit workflow (Figure 2A) is as follows:

**Figure 2.**
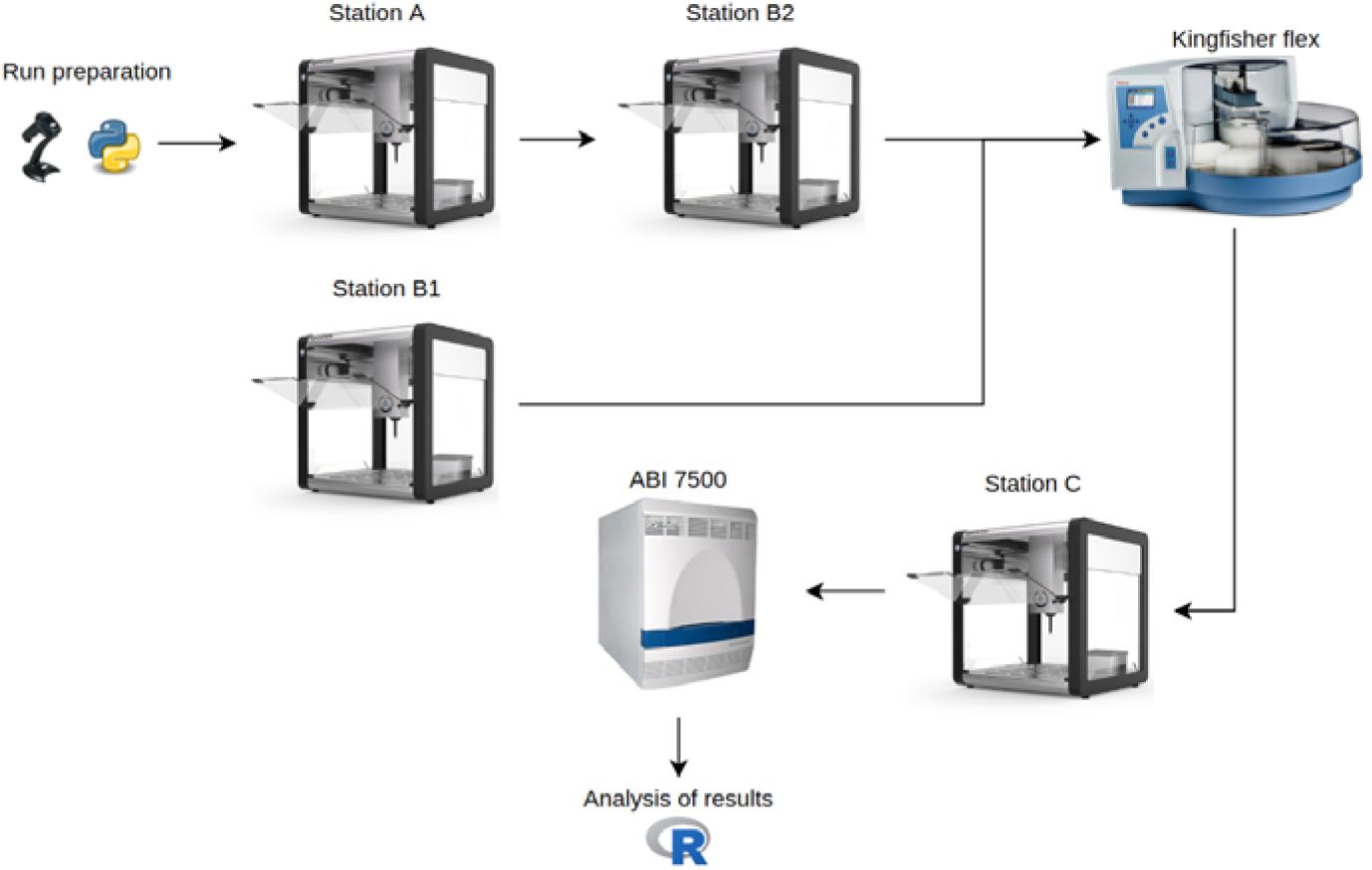
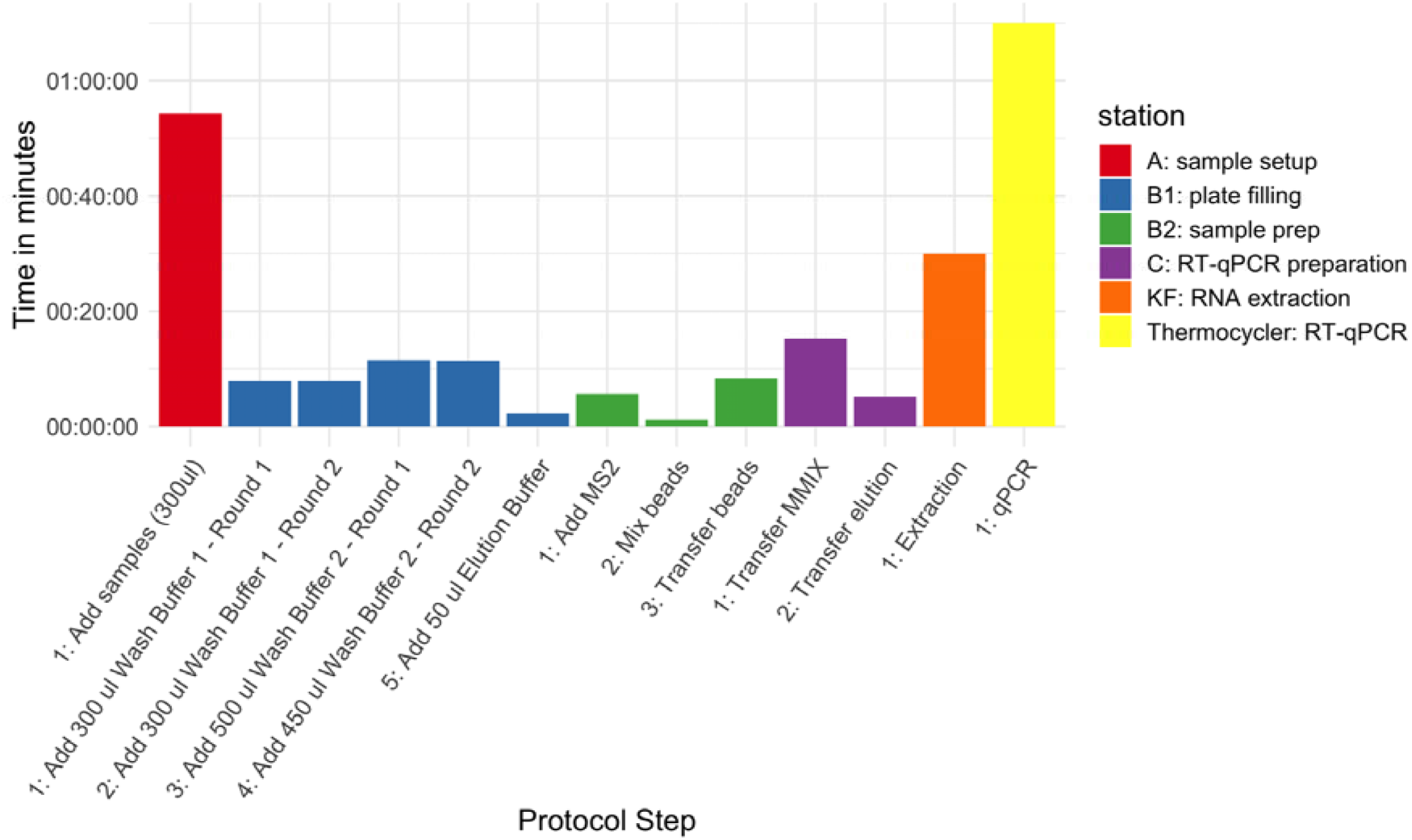
**A)** Diagram of the OT2-KF circuit. The full configuration of labware for OT2 stations is available as supplementary material. **B)** Run times in minutes for the different steps classified by station in the Kingfisher pathogen circuit using the MagMax™ Pathogen RNA/DNA kit.

1. Initial sample setup (OT-2 station A), 2. sample preparation (OT-2 station B1), 3. plate filling for Kingfisher Flex (OT-2 station B2), 4. RNA extraction step (Kingfisher Flex station), 5. RT-qPCR preparation (OT-2 station C), and the 6. RT-qPCR (ABI 7500 Fast thermocycler).

Each OT-2 station can have two pipettes simultaneously installed. The OT-2 pipettes have three different capacities (20, 300, 1000 µl) and can be either single-channel or multi-channel. Letters in the OT-2 station nomenclature are defined by a combination of such pipettes, therefore stations B1 and B2 can be run on the same robot. A complete description of every station with the labware needed is available as supplementary material

The complete process for testing 96 samples takes about 4h (Table 1) and it can be operated by a single laboratory technician. The longest step was the RT-qPCR amplification and detection using the ABI 7500 Fast thermocycler. Theoretically, a new run could start every 70 minutes; however the inactivation of samples becomes the real bottleneck. A summary of the steps carried out in each station and their executing time is indicated in Figure 2B.

**Table 1.** Number of steps and running times in minutes for a full run with 96 samples across the different stations. Station A and B2 can run simultaneously.

| Station | Steps | Total time (minutes) |
| --- | --- | --- |
| A: sample setup | 1 | 54 |
| B1: plate filling | 5 | 41 |
| B2: sample prep | 3 | 59 |
| C: RT-qPCR preparation | 2 | 20 |
| KF: extraction | 1 | 30 |
| Thermocycler: RT-qPCR | 1 | 70 |
| **Total** |  | 234 |

### Comparison of OT-2-KF pathogen circuit with other platforms

The European Molecular Genetics Quality Network (EMQN) is a provider of External Quality Assessment (EQA) services that are essential for any laboratory seeking to provide a quality service. The QCMD 2020 Coronavirus Outbreak Preparedness EQA Pilot Study aims to assess the proficiency of laboratories in the detection of different coronavirus genotypes including the new variant SARS-CoV-2. Eight inactivated samples were analysed with four different platforms; Roche, Cobas 6800, Hamilton-Seegene, and KF pathogen circuit (see supplementary methods).

We detected five out of eight samples as positive and the others as negative, with consistent results across the four platforms, even though the target regions were different (Table 2a). The internal control was correctly amplified for all samples and platforms.

**Table 2.**
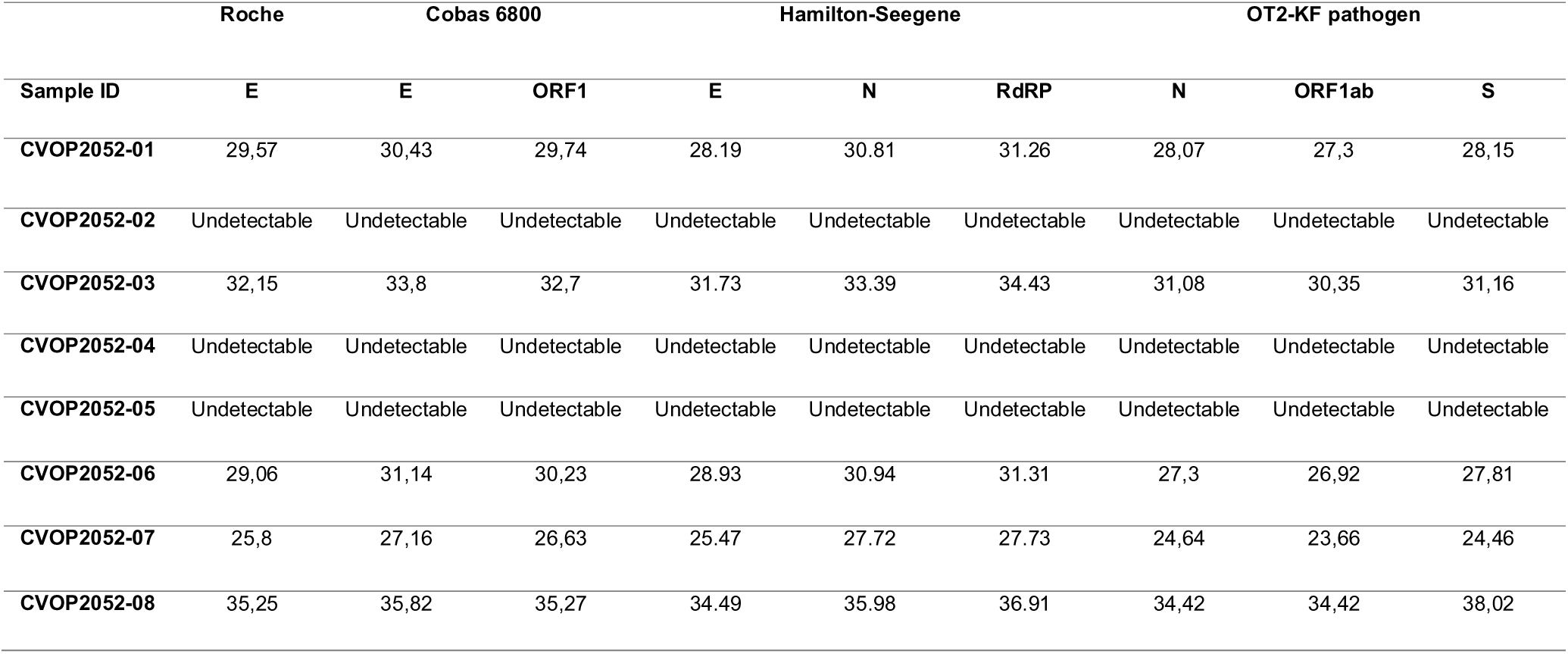
Comparison of four different platforms in the detection of SARS-CoV2 from external quality samples, including our circuit OT2-KF. Every platform has its own targets and primers.

Ct levels are inversely proportional to the amount of target nucleic acid in a sample. The range of Ct for Roche platform is 25.8-35.25, 25.47-35.82 for Cobas 6800, and 27.72-36.91 for Hamilton-Seegene. The range of Ct for the platform OT2-KF pathogen was 23.66 - 38.02, with consistently lower values except for the S gene in sample CVOP2052-08.

After submitting our results to the EMQN, we received the original content of each sample tested (Table 2b). We identified correctly all the SARS-CoV-2 strains. The other strains of coronavirus were not detected, showing that the tests have specificity for SARS-CoV-2.

**Table 2b.** Sample content of the EQA Pilot Study. *Sample code*: QCMD panel sample codes for the samples distributed to laboratories. *Sample content*: Content of the individual panel samples and, where applicable, the subtype or stain of the pathogen. *Expected result*: Expected outcome when testing for SARS-CoV-2. *Log10 dPCR Copies/ml*: The value obtained using a digital droplet PCR assay (modified from Corman et al 14). Samples CVOP20S2-07, 01, 03, 08, are in a calibrated dilution series. CVOP20S2-06 is a duplicate sample of CVOP20S2-01.

| Sample code | Sample content | Expected result | Log10 dPCR Copies/ml |
| --- | --- | --- | --- |
| CVOP20S2-01 | Novel Coronavirus SARS-CoV-2 | Positive | 4.30 |
| CVOP20S2-02 | Coronavirus NL63 | Negative | 4.64 |
| CVOP20S2-03 | Novel Coronavirus SARS-CoV-2 | Positive | 3.30 |
| CVOP20S2-04 | Coronavirus OC43 | Negative | 4.03 |
| CVOP20S2-05 | CV Negative | Negative | - |
| CVOP20S2-06 | Novel Coronavirus SARS-CoV-2 | Positive | 4.30 |
| CVOP20S2-07 | Novel Coronavirus SARS-CoV-2 | Positive | 5.30 |
| CVOP20S2-08 | Novel Coronavirus SARS-CoV-2 | Borderline Positive | 2.30 |

### Comparison of OT-2-KF pathogen circuit with OT2-KF viral pathogen II

Fifteen previously frozen samples with known outcome were run across the two KF circuits in two separate batches. All of the samples tested had qualitatively the same results between kits (Table 3). Sample 200905815 had one amplified target (N – nucleocapsid gene) with a very high Ct (37.9), compatible with a patient after infection with some remains of the virus.

**Table 3.** Comparison of fifteen samples using the two automated circuits implemented in our lab using Thermo Fisher kits: OT2-KF pathogen and OT2-KF Viral Pathogen II.

| Sample | Kit Pathogen |  |  |  | Kit Viral Pathogen II |  |  |  |
| --- | --- | --- | --- | --- | --- | --- | --- | --- |
|  | N | ORF | S | MS2 | N | ORF | S | MS2 |
| 202228221 | Undetectable | Undetectable | Undetectable | 26,4 | Undetectable | Undetectable | Undetectable | 26,5 |
| 200907564 | 22,9 | 23,1 | 22,7 | 22,9 | 23,9 | 23,4 | 22,8 | 27,1 |
| 200907556 | 27,3 | 27,9 | 27,8 | 26,7 | 29,2 | 29,2 | 28,7 | 27,5 |
| 200907560 | 31,4 | 31,3 | 31,6 | 26,4 | 32,5 | 34,5 | 33,0 | 27,4 |
| 200904740 | 28,9 | 30 | 29 | 26,7 | 32,7 | 34,7 | 34,1 | 25,6 |
| 200904403 | Undetectable | Undetectable | Undetectable | 26,3 | Undetectable | Undetectable | Undetectable | 27,5 |
| 200905798 | 31,9 | 31,4 | 29,7 | 26,8 | 33,4 | 33,3 | 32,3 | 28,7 |
| 200905815 | Undetectable | Undetectable | Undetectable | 25,9 | 37,9 | Undetectable | Undetectable | 28,2 |
| 200907593 | 17,9 | 18,5 | 18,6 | 26,3 | 20,0 | 20,2 | 19,8 | 28,0 |
| 200907603 | 26,3 | 26,8 | 24,8 | 26,7 | 27,0 | 27,2 | 26,9 | 27,0 |
| 200907154 | 21,8 | 21,9 | 21,9 | 26,9 | 22,5 | 22,7 | 22,3 | 25,1 |
| 200906987 | 25,7 | 26,2 | 24,9 | 27,3 | 26,9 | 26,8 | 26,2 | 27,8 |
| 202230162 | 23,4 | 23,4 | 23,2 | 26,5 | 25,0 | 25,1 | 24,8 | 27,1 |
| 200903314 | 27,8 | 27,9 | 27,5 | 25,7 | 29,3 | 28,7 | 28,4 | 26,8 |
| 202224392 | Undetectable | Undetectable | Undetectable | 25,9 | Undetectable | Undetectable | Undetectable | 27,0 |

The Ct values for the OT-2-KF viral pathogen II were 1.6 higher on average. One possible explanation for this small difference is that the samples had an additional round of freezing-unfreezing as they were processed in the second batch.

## DISCUSSION

### Working framework

We successfully created a framework to develop RT-qPCR automated tests OT-2 robots and additional commercial systems, validating it up to production level and establishing a working circuit in our centre. According to our EMQN validation, the results obtained with this platform are equivalent to other commercial solutions.

The framework includes classes and functions that aim to facilitate the use of the liquid handler robots. For example, by creating a new class for liquids, we have been able to add a functionality that matches other liquid handler robots available on the market. All attributes that depend exclusively on liquid properties are described and can be modified in the object definition.

An advantage of establishing circuits based on independent modules is that some processes can run in parallel, resulting in an increased turnaround capacity of testing during a whole working day. Taking into account the station times in the OT-2-KF circuit (Figure 1), the longest step is the RT-qPCR amplification and detection using the ABI 7500 Fast thermocycler. Therefore, theoretically one run could start every 70 minutes. However, the real bottleneck is the inactivation of samples, which is a non-automated procedure that requires of specialized facilities and microbiology laboratory technicians.

### OT-2 advantages and limitations

The OT-2 stations are a promising solution to increase the SARS-CoV-2 testing capability since it is an affordable open source platform for liquid handling. Open-source platforms have advantages compared with other platforms such as cost-effectiveness, flexibility and fast adaptation to laboratory needs. Open-source platforms are as strong as their community. Working with them can be challenging at first but every actor should greatly benefit as the community of users and developers grow, testing and improving existing code and developing new functions and capabilities for the platform. Albeit growing, the OT-2 community is still small and there is not much code available. In that sense, we contribute with code to mimic capabilities available in other platforms, as mentioned above.

However, there are some capabilities that are difficult or impossible to do in OT-2 stations. For example it lacks hardware to control if a sample was correctly aspirated or detect clot formations. The RNA extraction is the most challenging step within the circuit. It can be theoretically done within an OT-2 using a magnetic module and we are currently testing different protocols with partial success to our standards. In the circuits presented here, the extraction is done in a separate robot, the KingFisher™ instrument which incorporates an agitator.

Sample tracking is also essential to ensure the quality of the diagnosis in a clinical setting. Currently, there is no hardware or software in the OT-2 robots to support it and we rely on external barcode readers for individual samples and fixed well positioning, with a series of custom automation scripts.

To setup a circuit like this from scratch it is necessary to have both in-house programming and molecular biology expertise. Even if templates and working code is available it will still be necessary to make changes, adjustments, and understand the biological context. In that sense, is important the figure of bioinformatics professionals that are being slowly incorporated in diagnostic environments such as hospitals.

### Hit the ground running

Although the first pandemic peak has been contained, increasing RT-qPCR capacities in laboratories should not leave the agenda, in preparedness for possible upcoming waves. According to the World Health Organization (WHO), diagnostic laboratories are one of the fundamental pillars for the acute response against the pandemic: “Countries should prepare laboratory capacity to manage large-scale testing for COVID-19 domestically, through public, private and academic laboratories.”^13^.

These strategies will only be possible if we develop a high capacity to perform RT-qPCR tests. In this sense, the platform discussed here is versatile, affordable, and can be re-purposed for other tasks. Circuits built with this technology can be useful to increase the testing capacity of established laboratories and provide freedom during acute shortages. They can also be an optimal solution for testing labs that currently do not have any automated platform in place.

## AVAILABILITY

Code, documentation, and station diagram and descriptions are available in the CDB GitHub repository: https://github.com/CDB-coreBM/covid19clinic

Codes for Kingfisher robot are available in Thermo Fisher’s web site:

Pathogen. https://www.thermofisher.com/order/catalog/product/4462359#/4462359

Viral Pathogen II. https://www.thermofisher.com/order/catalog/product/A48383#/A48383

## SUPPLEMENTARY DATA

Supplementary methods and material is available online.

## AUTHOR CONTRIBUTIONS

JLVC, AGU, and EGR conceived the system, wrote code, and performed the robot testing; ETR, MJMY, AVG, F.M.L carried out experiments and provided samples and advise; JAPB helped supervise the project; JLVC wrote the paper with input from all authors.

## ACKNOWLEDGEMENT

We thank M.D. Jiménez, Anabel Martínez, Mar López, Marta Parera, Núria Palau, Víctor Pastor, and Paula Sánchez for laboratory support and advice. Elena Roel for insightful comments. Ojas Patel for technical support. The authors thank the NGO COVIDWarriors for donating the Opentrons OT-2 stations to the Hospital Clinic of Barcelona (Spain).

