## Supplementary material 1 for "ROBOCOV: An affordable open-source robotic platform for SARS-CoV-2 testing by RT-qPCR"

CDB

Centre de Diagnòstic Biomèdic

### OT2-KF station configurations

CLÍNIC
BARCELONA
Hospital Universitari

### Summary



This supplementary material contains the pipette configuration, materials and disposition for the stations included in the OT2-KF Pathogen circuit and the OT2-KF Viral Pathogen II (VPII) circuit.

OT2-KF Pathogen and Viral Pathogen II share stations A and C labwares and only differ in stations B1 and B2.

The total material needed for either circuit is at the end of the supplementary material.

CLÍNIC BARCELONA Hospital Universitari

Centre de Diagnòstic Biomèdic CDB

### OT2-KF Station A

Sample preparation

# KF-A

### Disposition

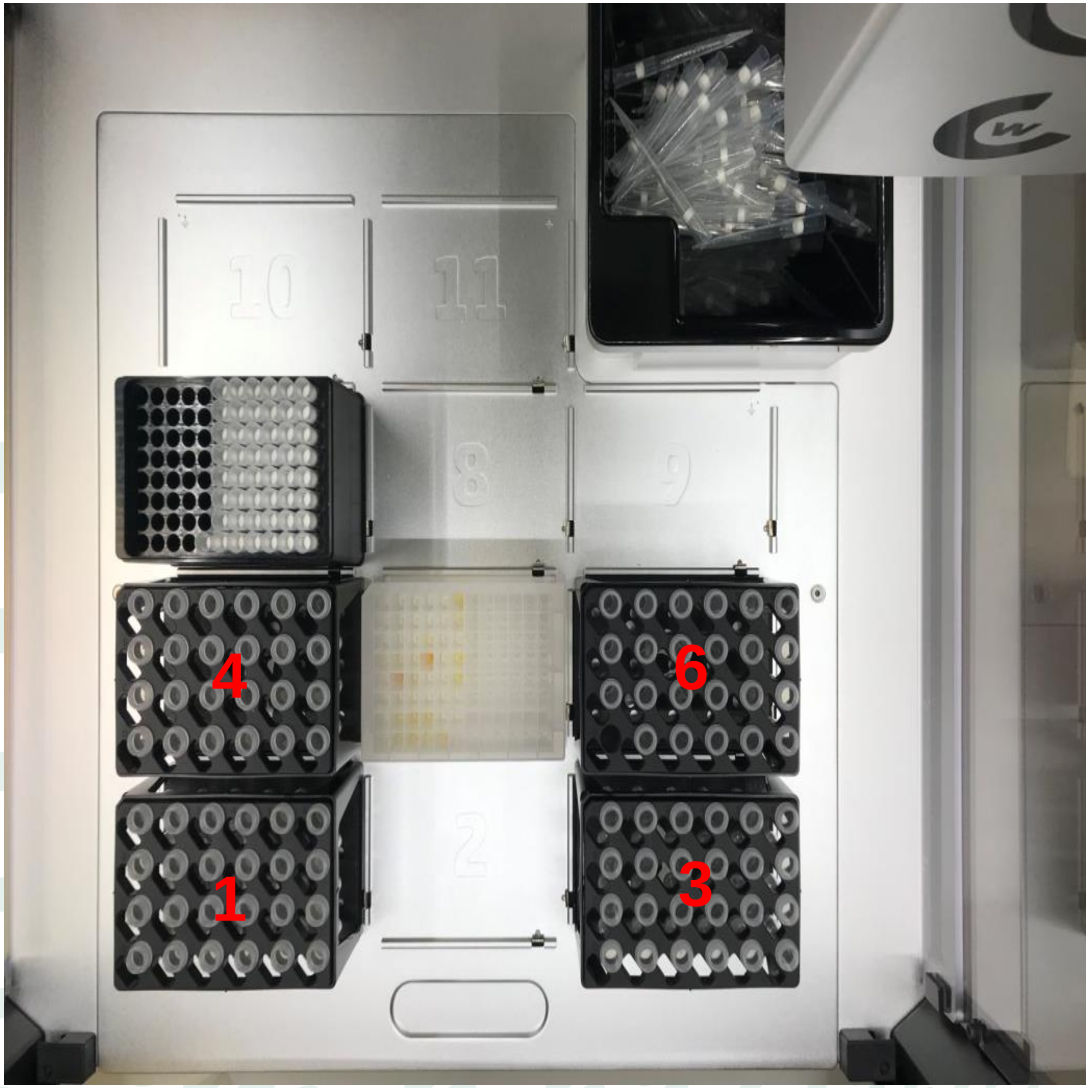


**Pipettes**

*Left:* p1000

*Right:* p20

**Slots**

1. Samples-1 (2ml)
2. Tiprack 20
3. Samples-2 (2ml)
4. Samples-3 (2ml)
5. KingFisher Deepwell plate
6. Samples-4 (2ml)
7. Tiprack P1000
8. *Empty*
9. *Empty*
10. *Empty*
11. *Empty*
12. Wastebin

CLÍNIC BARCELONA Hospital Universitari

Centre de Diagnòstic Biomèdic CDB

### OT2-KF Pathogen Station B1

Reagent setup
(plate filling)

### KF-B1 Disposition pathogen

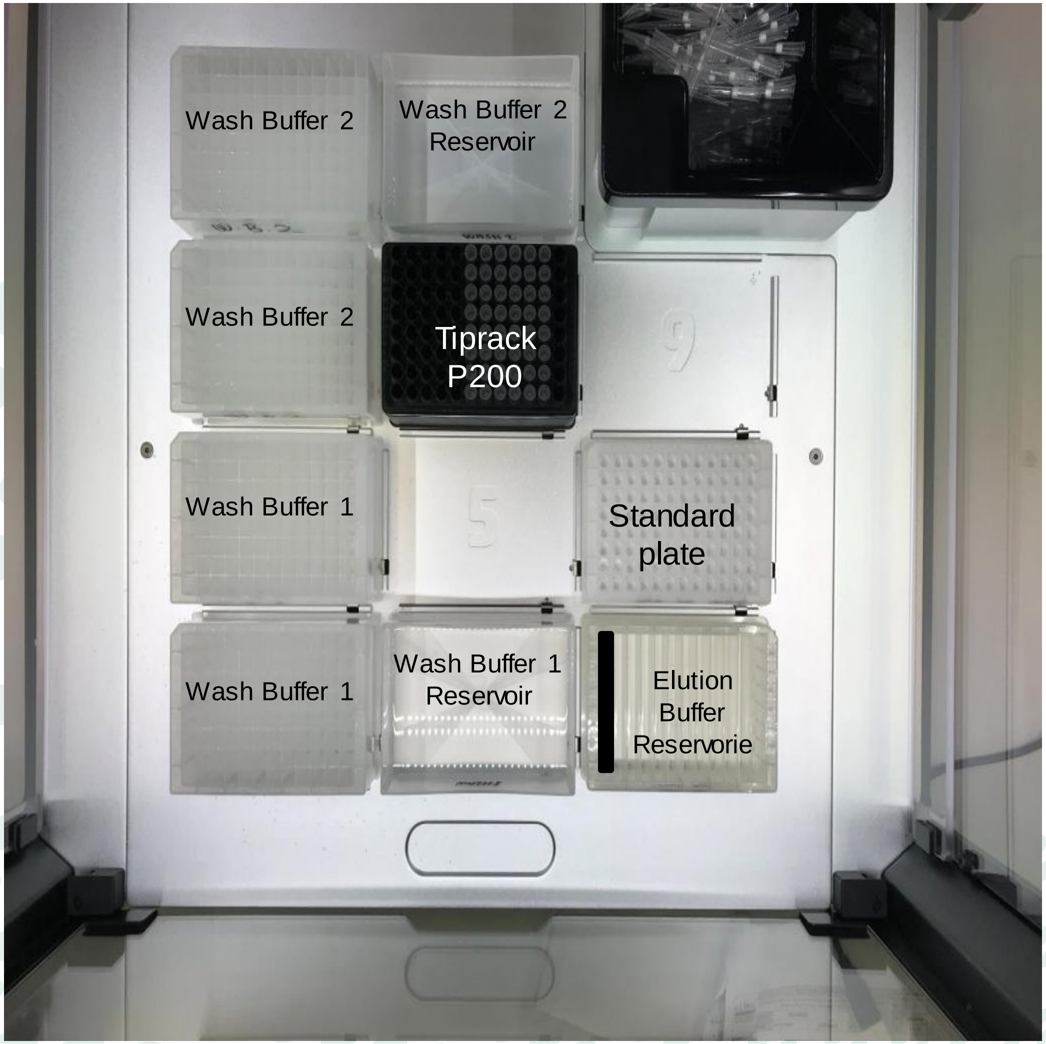


**Pipettes**
*Left:* m20 (not used)
*Right:* m300

**Slots**

1. KingFisher DeepWell plate
2. 300ml NEST reservoir
3. 12 channel NEST reservoir
4. KingFisher DeepWell plate
5. *Empty*
6. KingFisher Standard plate
7. KingFisher DeepWell plate
8. Tiprack 200
9. *Empty*
10. KingFisher DeepWell plate
11. 300ml NEST reservoir
12. Wastebin

### OT2-KF Pathogen Station B2

#### Sample preparation

KF-B2

### Disposition pathogen

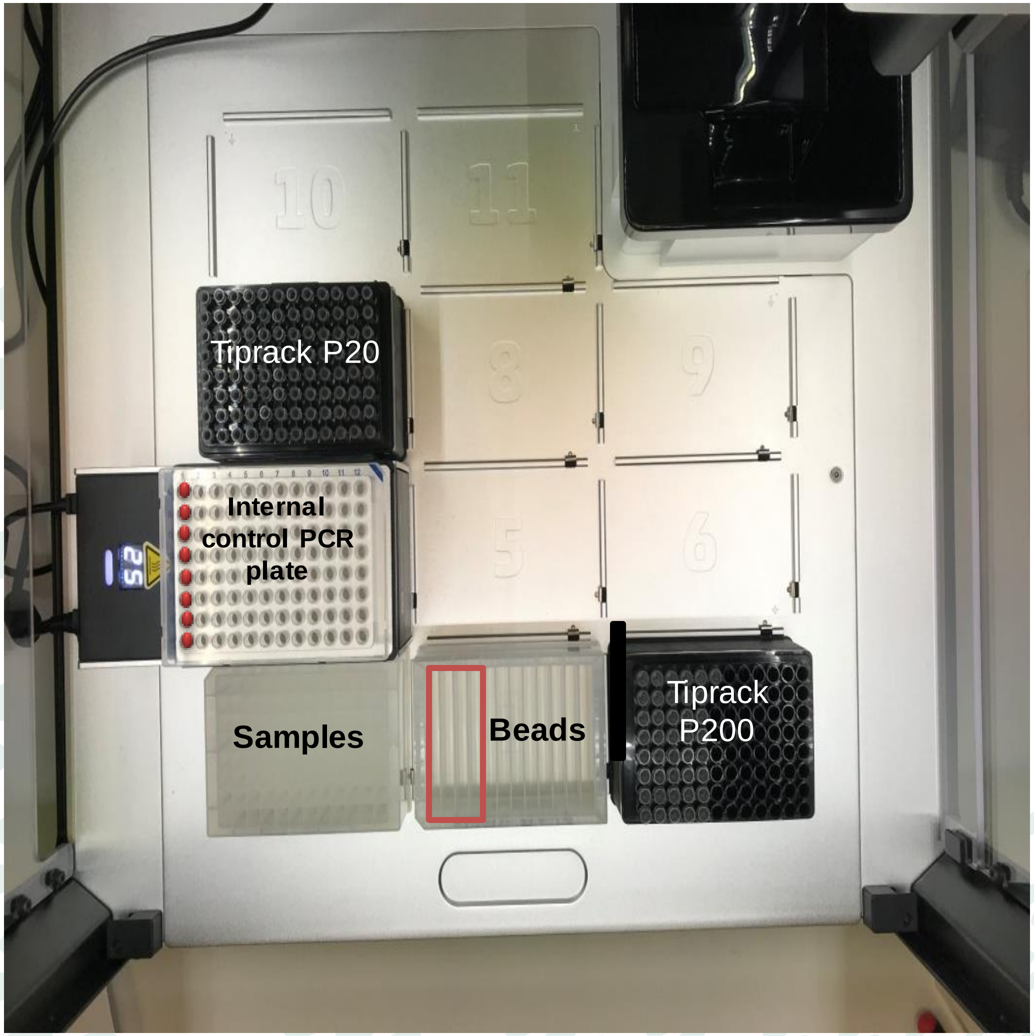


**Pipettes**
*Left:* m20
*Right:* m300

**Slots**

1. KingFisher DeepWell plate
2. 12 channel NEST reservoir (up to 4 channels for beads)
3. Tiprack 200
4. PCR plate + mod Temp.
5. *Empty*
6. *Empty*
7. Tiprack 20
8. *Empty*
9. *Empty*
10. *Empty*
11. Waste pool reservoir
12. Wastebin

### OT2-KF VPII Station B1

Reagent setup
(plate filling)

KF-B1

### Disposition VPII

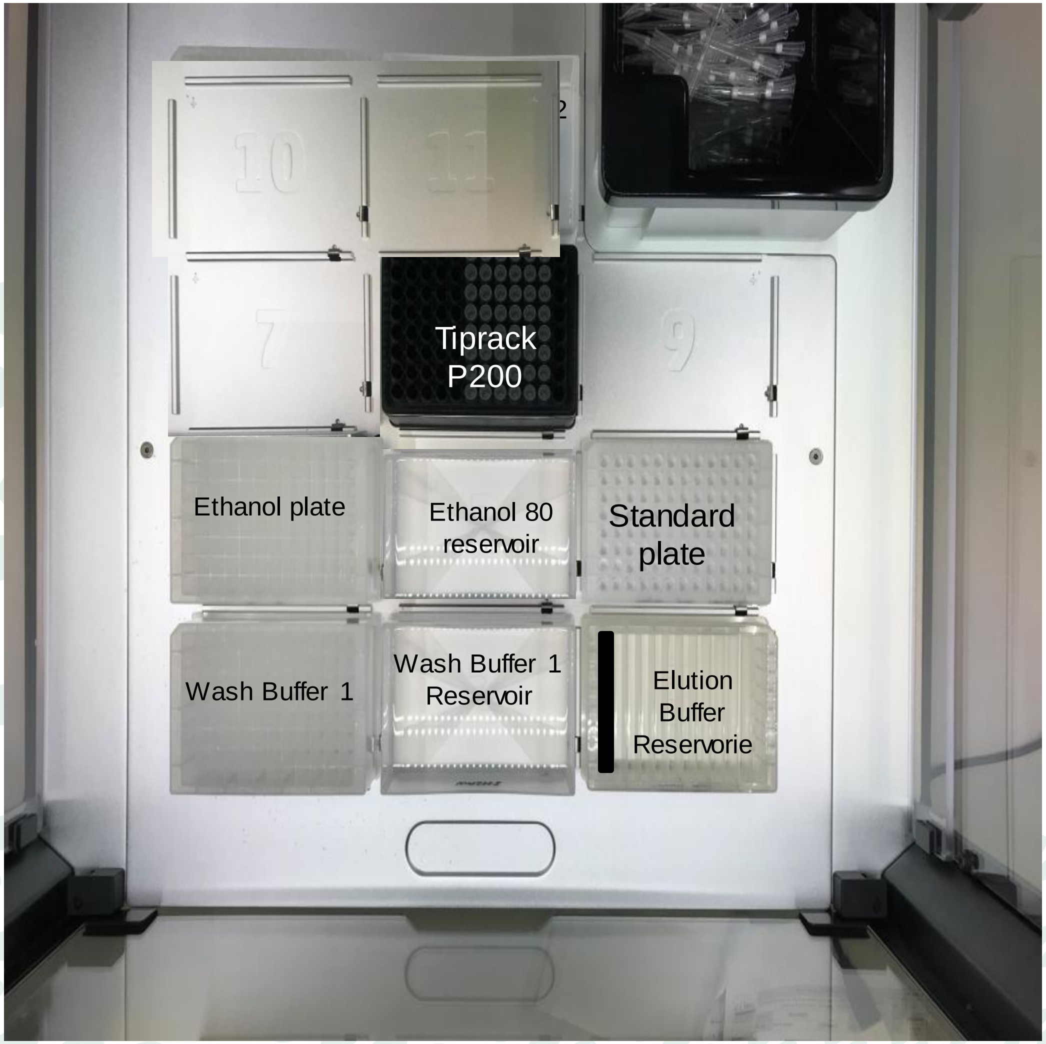


**Pipettes**

*Left:* m20 (not used)

*Right:* m300

**Slots**

1. KingFisher DeepWell plate
2. 300ml NEST reservoir
3. 12 channel NEST reservoir
4. KingFisher DeepWell plate
5. 300ml NEST reservoir
6. KingFisher Standard plate
7. *Empty*
8. Tiprack 200
9. *Empty*
10. *Empty*
11. *Empty*
12. Wastebin

CLÍNIC BARCELONA Hospital Universitari

Centre de Diagnòstic Biomèdic CDB

### OT2-KF VPII Station B2

Sample preparation

KF-B2

### Disposition VPII

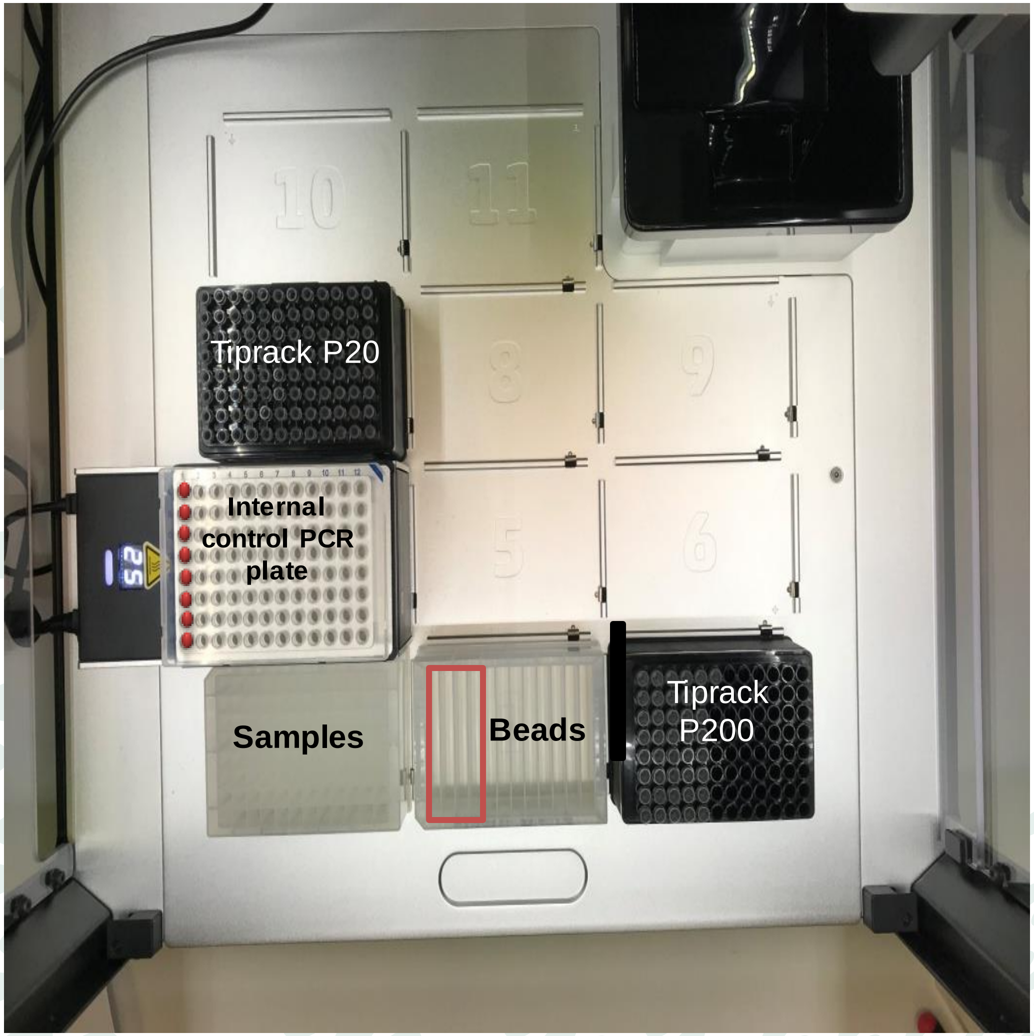


**Pipettes**

*Left:* m20

*Right:* m300

**Slots**

1. KingFisher DeepWell plate
2. 12 channel PerkinElmer 21mL reservoir
3. Tiprack 200
4. PCR plate + mod Temp.
5. *Empty*
6. *Empty*
7. Tiprack 20
8. *Empty*
9. *Empty*
10. *Empty*
11. *Empty*
12. Wastebin

Hospital Universitari

# OT2-KF
### Station C

qPCR
setup

KF-C

### Disposition

Centre de Diagnòstic Biomèdic
CDB

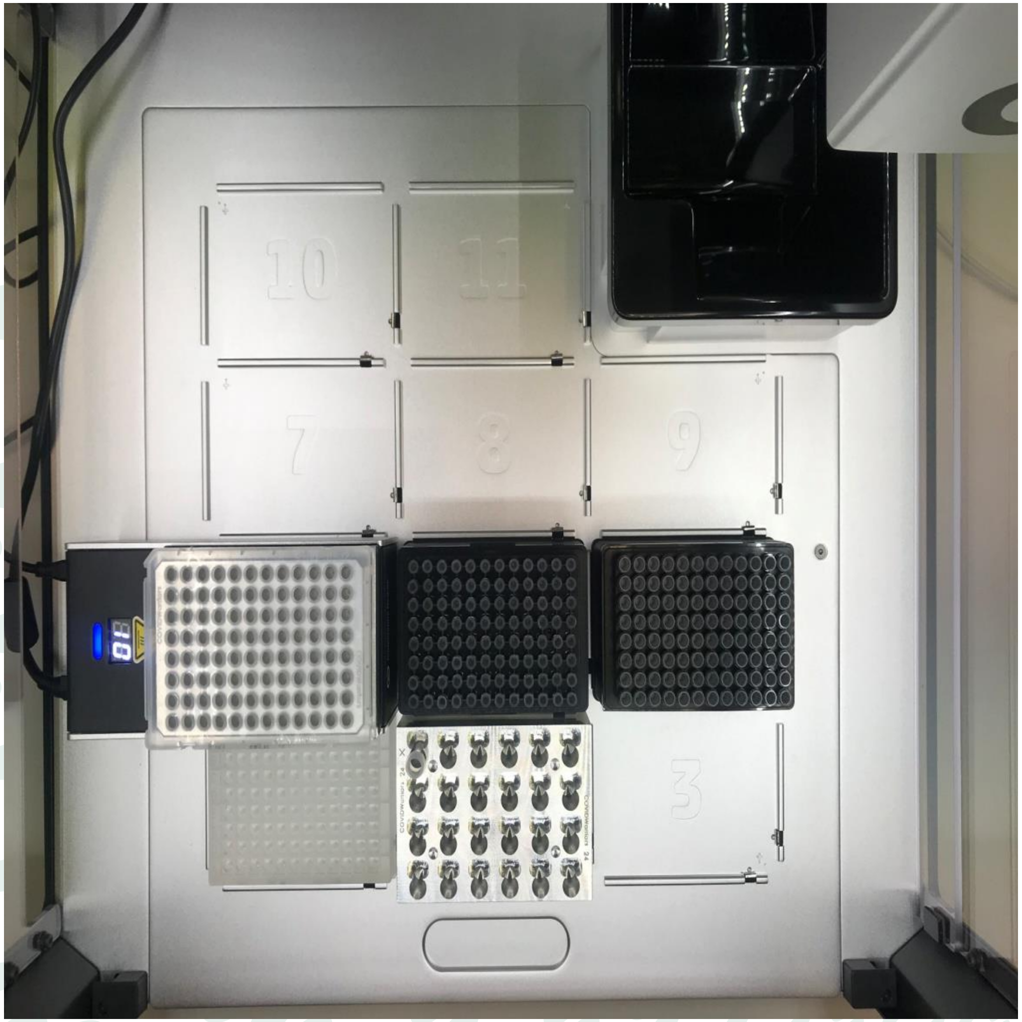


**Pipettes**

*Left:* p300

*Right:* m20

**Slots**

1. Deepwell
2. Mastermix (2 screwcaps, ref)
3. *Empty*
4. qPCR plate + mod Temp.
5. Tiprack 20
6. Tiprack 200
7. *Empty*
8. *Empty*
9. *Empty*
10. *Empty*
11. *Empty*
12. Wastebin

### Total material for one run

**Reusable**

2 Aluminium blocks 96
1 Aluminium block 24
4 racks with eppendorf adaptors
2 Temperature modules

**Consumable**

1 Tiprack 1000
3 Tiprack 200
2 Tiprack 20
5 KingFisher Deepwell plate
1 Standard KingFisher plate
1 PCR plate 96
1 qPCR plate 96
2 Reservoir 300 mL
2 Reservoir 12 channel NEST

### Total material for one run

**Reusable**

2 Aluminium blocks 96
1 Aluminium block 24
4 racks with eppendorf adaptors
2 Temperature modules

**Consumable**

1 Tiprack 1000

3 Tiprack 200

2 Tiprack 20

3 KingFisher Deepwell plate

1 Standard KingFisher plate

1 PCR plate 96

1 qPCR plate 96

2 Reservoir 300 mL

1 Reservoir 12 channel NEST

1 Reservoir 12 channel perkinelmer (21ml)
