## Supplemental methods for "ROBOCOV: An affordable open-source robotic platform for SARS-CoV-2 testing by RT-qPCR"

**EMQN quality controls**

The European Molecular Genetics Quality Network (EMQN) is a provider of External Quality Assessment (EQA) services that are essential for any laboratory seeking to maintain and provide a quality service. The QCMD 2020 Coronavirus Outbreak Preparedness EQA Pilot Study aims to assess the proficiency of laboratories in the detection of different coronavirus genotypes including the new variant SARS-CoV-2. Eight inactivated samples were analysed with four different platforms, including the workflow OT2-KF. Samples were mixed 1:1 with cobas omni Lysis Reagent (Roche) before being processed to allow comparison.

**rRT-PCR using Roche Modular Kit**

An automatic nucleic acid extraction from 400 µl of inactivated samples was performed on a MagNa Pure Compact instrument (Roche Applied Science, Mannheim, Germany) using the MagNA Pure Compact Nucleic Acid Isolation Kit I - Large Volume (Roche), following the manufacturer’s instructions. Equine arteritis virus (EAV), a positive-sense single-stranded RNA virus, was added to all samples prior to RNA extraction and served as an internal extraction and amplification control. For detection of SARS-CoV-2, a 76 bp long fragment from the E gene was amplified with specific primers and detected with a FAM label hydrolysis probe using a LightMix Modular kit (Roche). The assay detects SARS and 2019-nCoV pneumonia virus (bat-associated SARS related Sarbecovirus). RT-PCR was performed in combination with the LightCycler® Multiplex RNA Virus Master (Roche) on the LightCycler 480 Real-Time PCR System (Roche) following recommended cycling conditions: reverse transcription at 55 °C for 3 min, and 95 °C for 20 sec, followed by 45 cycles of PCR at 95 °C for 3 sec and 60 °C for 30 sec (Corman).

**rRT-PCR using COBAS 6800**

Cobas 6800 (Roche) is a fully automated sample-to-result platform, including the sample supply module, the transfer module, the processing module, and the analytic module. For detection of SARS-CoV-2, a two-target RT-PCR is used: one targeting ORF1, a non-structural region that is unique to SARS-CoV-2 (target 1), and the second targeting a conserved region in the structural protein envelope E gene for pan-Sarbecovirus detection (target 2). The pan-Sarbecovirus primers and probe should also detect the SARS-CoV-2 virus. The test utilizes RNA internal control for sample preparation and PCR amplification process control. Automated data management was performed by the manufacturer’s software, which assigns test results for all tests. In our study, 400 µl of aliquots with lysis buffer were extracted on the Cobas 6800 system and tested following the manufacturer’s instructions. Testing was performed in batches of 94 samples plus one negative and positive control each.

**rRT-PCR using SeeGene**

RNA was extracted using Seegene STARMag 96 X 4 Universal Cartridge Kit (Seegene) on the Microlab STAR Liquid Handling System (Hamilton). rRT-PCR protocol with the Allplex™ 2019-nCoV Assay was automated: prepared on the Microlab STAR Liquid Handling System (Hamilton) and detected on the CFX96 Touch Deep Well Real-Time PCR Detection System (Bio-Rad). C_t_ from FAM (E gene), Cal Red 610 (RDRP gene), Quasar 670 (N gene) and HEX (internal control) were acquired.

**Interpretation of results**

A sample was considered negative if the internal control was amplified but not the viral genes. A specimen was considered invalid when there was no amplification of the internal control. Samples were considered positive when a signal was detected at C_t_<38 for Roche (Modular and COBAS 6800) or Ct<37 for any gene in case of Seegene technique.
